# Characterizing the Cellulose Binding Interactions of Type-A Carbohydrate-Binding Modules using Acoustic Force Spectroscopy

**DOI:** 10.1101/2023.10.18.563009

**Authors:** Gokulnath Ganesan, Markus Hackl, Shishir P.S. Chundawat

## Abstract

A critical understanding of carbohydrate binding modules (CBMs) is vital for the manipulation of a variety of biological functions they support, including biomass deconstruction, polysaccharide biosynthesis, pathogen defence, and plant development. The unbinding characteristics of CBMs from a polysaccharide substrate surface can be studied using rupture force measurements since it enables a quantitative inference of binding properties through the application of dynamic force spectroscopy (DFS) theory. With the increase in usage of CBMs for diverse applications, it is important to engineer and characterize CBMs that have desired sets of interactions with various carbohydrate substrates. However, though the effect of mutations in the binding motif residues is known to influence CBM binding affinity, its effect on the rupture forces is still not well quantified. This is primarily due to the low experimental throughput of most single-molecule DFS techniques available to characterize the force-induced dissociation of protein-ligand interactions. Here, we have determined the rupture forces of microscopic beads functionalized with various wild-type and mutant CBMs using a highly multiplexed DFS technique called Acoustic Force Spectroscopy (AFS). We have characterized the acoustic force-induced dissociation of specifically two type A CBMs (i.e., CBM3a and CBM64) and relevant seven binding motif targeting CBM mutants unbinding from a nanocellulose surface, over a broad range of DFS loading rates (i.e., 0.1 pN/s to 100 pN/s). Our analysis of the rupture force DFS data yields apparent CBM-cellulose bond interaction parameters, which enables a quantitative comparison of the effect of corresponding mutations on cellulose-CBM binding interactions that compares favorably with results from classical bulk ensemble based binding assays. In summary, detailed insights into the rupturing mechanism of multi-CBM fused domains provides motivation for usage of specific constructs for industrial biotechnological applications.

## Introduction

Carbohydrate binding modules (CBMs) are non-catalytic auxiliary domains that bind to specific polysaccharides and bring the tethered catalytic domain in close proximity to the substrate. Due to the key role they play in helping enzymes associate with their substrates, CBMs have diverse biotechnological applications ranging from transgenic crop engineering, paper pulp processing, analytical detection of carbohydrates, biomass deconstruction, and development of novel biomaterials [1], [2]. CBMs are also an excellent model system for studying protein-carbohydrate recognition due to the broad range of ligand specificities that these proteins can exhibit [3]–[5]. One of the widely studied applications of CBMs is in enabling the biocatalytic deconstruction of biomass, since the nature of interaction between CBMs and cellulose has a strong impact on the activity of cellulolytic enzymes [6], [7]. CBMs with moderate binding affinity are known to help in achieving higher enzymatic activity due to an optimum trade-off between substrate specificity and non-productive binding [8], [9]. Previous studies have also evolved more efficient enzymes for deconstructing lignocellulosic biomass by altering CBM-cellulose interactions through point mutations on the CBM binding motif sites [10], [11].

The protein-substrate interaction can be characterized using classical solid-state depletion assays [12] or more elaborate techniques such as quartz crystal microbalance (QCM) [11], [13] and fluorescence recovery after photobleaching (FRAP) [14]. All such classical methods rely on the equilibrium binding and unbinding kinetics analysis under simple conditions. However, here highly multivalent protein-substrate interactions are often classified by such simple methods as irreversibly bound proteins providing limited mechanistic understanding of more complex CBM-carbohydrate binding phenomenon [15]. To overcome this limitation, we have recently developed a DFS method to study the force-induced dissociation of proteins from model cellulose surfaces [16]. Here, we further demonstrate the use of our previously developed protocol by systematically characterizing the ce Here, we have determined the rupture forces of microscopic beads functionalized with various mutant CBMs using a highly multiplexed DFS technique called Acoustic Force Spectroscopy. Llulose surface unbinding forces of two distinct type A CBMs, namely CBM3a from *Clostridium thermocellum* and CBM64 from *Spirochaeta thermophila*. In addition to the wild-type proteins, we have characterized the unbinding behaviour of the alanine point mutations of critical binding motif residues known to be important for CBM-cellulose binding interactions. The mutations at different binding sites of the mutant CBMs, as represented in **Figure 1**A, could possibly result in distinct binding interactions with the cellulose surface, as previously reported [17], [18]. Characterizing the effect of these mutations using DFS is an important step toward engineering cellulases for more efficient deconstruction of cellulosic biomass into fermentable sugars.

**Figure 1:**
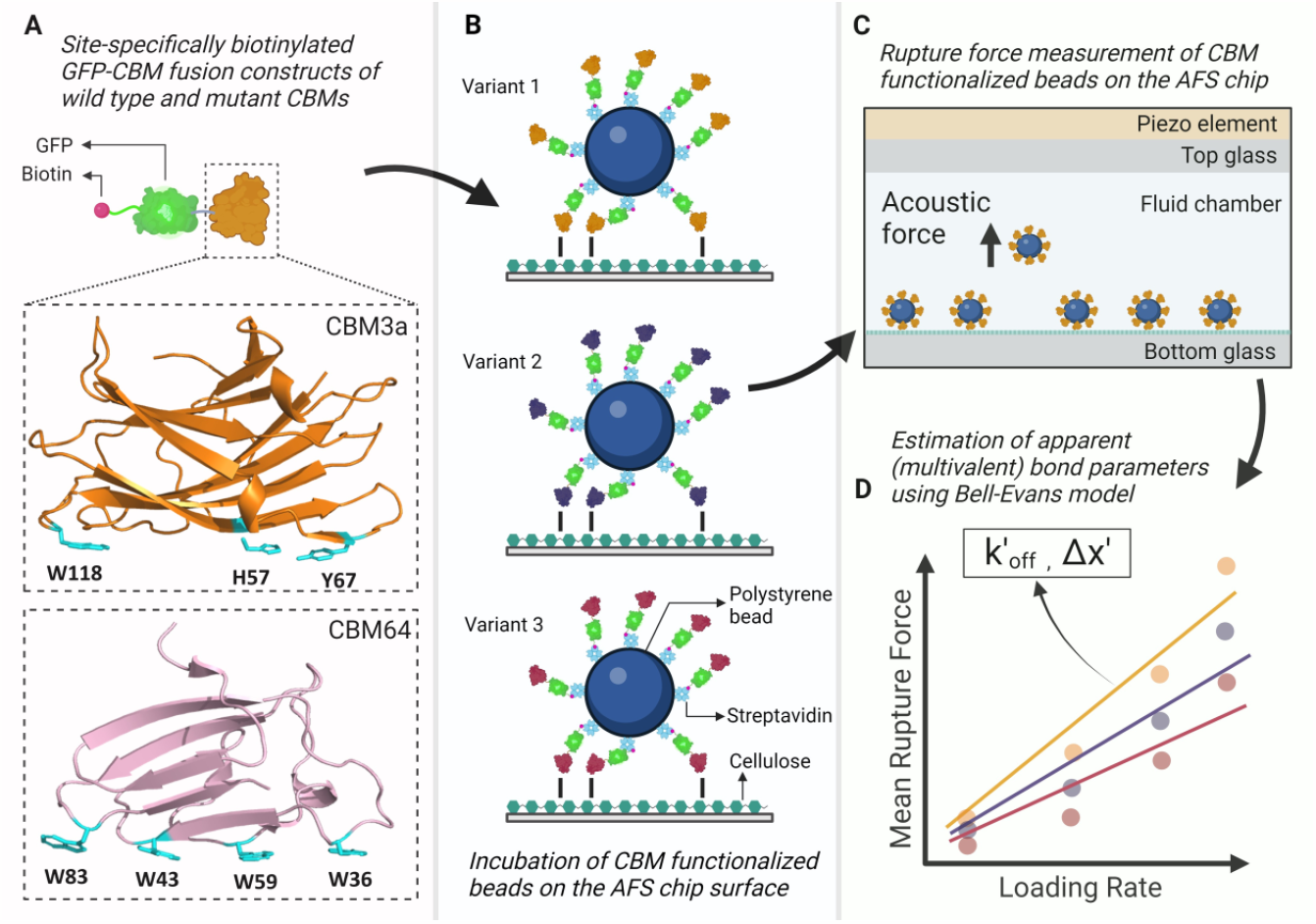
Overview of the workflow used to characterize the unbinding of the wild type and mutants of CBM3a and CBM64 from a cellulose surface using AFS. (A) By mutating the highlighted amino acids to Alanine one at a time, we obtained 3 mutants of CBM3a and 4 mutants of CBM64. A biotin-GFP-CBM construct was made following the procedure in [16]. (B) Streptavidin coated polystyrene beads are functionalized with the CBM constructs and allowed to bind onto the nanocellulose that was deposited in the fluid chamber of the AFS chip. Beads functionalized with three variants of CBM (wild type and mutants) and bound to the cellulose surface are depicted here. (C) A cross-sectional view of the AFS chip showing force-induced rupturing of a bead due to the upward acoustic force exerted on it. (D) The rupture force data of the beads are analyzed to obtain mean rupture forces at various loading rates of the applied force. Apparent (multivalent) bond parameters, k’_off_ and Δx’, of a CBM are obtained by fitting the Bell-Evans model to the mean rupture force data at multiple loading rates.

Since the initial protocol development [16], we have now further modified the protocol with minimal exposure of the chip surface to passivation proteins (BSA and Casein) with buffers used for surface passivation and bead preparation and further avoiding the usage of bleach. This results in minimal changes to the cellulose surface deposited on the AFS chip, allowing us to reliably screen multiple mutant CBMs over a much longer duration than previously reported. AFS technique has proven to be a reliable methodology to advance our understanding of DNA-protein, cell mechanics [19]–[22], and enzyme-substrate interactions, particularly for bioengineering applications [17]. The usage of AFS has enabled our study of the unbinding behaviour of polystyrene beads homogenously functionalized with mutant CBMs with a model nanocellulose surface over a wide range of unbinding forces and at high experimental throughput to enable rapid DFS data collection.

## Methods

### Chemical Reagents

Unless otherwise mentioned, all reagents were either purchased from VWR International, Fisher Scientific, or Sigma-Aldrich. Streptavidin-coated beads (nominal diameter 5.14 μm) were purchased from Spherotech, Inc. Sulfuric acid-hydrolyzed nanocellulose (NCC) was kindly donated by Richard Reiner, USDA Forest Product Laboratory, Madison, Wisconsin [23].

### Chip Preparation

The deposition of nanocrystalline cellulose (NCC) on the AFS chip was done following the protocol described in [17]. A buffer containing 10 mM phosphate buffer, supplemented with 2.5 mg/ml Bovine Serum Albumin (BSA) and Casein, respectively, and 5.6 mg/ml of Pluronic F-127 (herein referred to as B2) was used to passivate the nanocellulose surface once after the deposition and drying of nanocellulose on the AFS chip, as described in [17]. In addition, Pro-Clin 300 (0.05% v/v) was added to all the buffers to prevent microbial growth on the cellulose surface, since the chip was used up to 16 days. For short-term storage, the chip was rinsed with 200 μl DI water and the microfluidic chamber was filled with DI water supplemented with Pro-Clin 300 (0.05% v/v) and sealed with parafilm to prevent drying.

### Creation of Biotinylated Proteins

Based on the plasmids for CBM3a, CBM64 and their alanine mutants prepared in [11], we introduced the AviTag [24] between the existing Histidine tag and the GFP domain using sequence and ligation independent cloning (SLIC) as previously described [16]. All plasmid sequences were verified by DNA sequencing (Genscript, Piscataway NJ). The biotin was added to the AviTag *in vivo* and the proteins were purified as outlined previously [16].

### Bead Functionalization with CBMs

A 10 mM phosphate buffer solution with 0.95 μg/ml Pluronic F-127 was used during bead functionalization and washing (herein referred to as WB). The CBM constructs were diluted to the required concentrations in WB. In a PCR tube, 10 μl of this dilution was incubated with 5 μl of streptavidin-coated beads for 15 min in a rotisserie to functionalize the beads with the CBMs. Next, 100 μl of WB were added, and the sample was spun down for 3 min. The supernatant was removed while the settled beads are again resuspended in WB and washed once more. The addition of Pluronic F-127 in WB was necessary to allow the settling of the functionalized beads during the washing step. After the second washing step, the beads were resuspended in 10 mM phosphate buffer only.

For functionalizing the beads with two CBM mutants, 5 μl of a 500 nM dilution of each mutant CBM construct and 5 μl of the beads were mixed, incubated and washed following the above methodology.

### Determining the rupture forces of CBM functionalized beads

Stokes force calibration to correlate the applied power and force experienced by the bead was done following the protocol described in [16], except the cellulose surface was passivated only in B2 buffer for 15 minutes. The beads were allowed to settle on the surface for 5 minutes, following which a look-up-table (LUT) was created to determine the position of the bead perpendicular to the surface [25]. The beads were tracked with a moving region of interest (ROI) at a frame rate of 40 Hz and a 10x magnification objective. First, the beads are tracked in the absence of any force for 15 seconds to determine the anchor point. Then, a small force (0.5 pN – 1.5 pN) is applied for 1 second. This force was large enough to lift the bead to the acoustic pressure node (z-node), but small enough to ensure that enough data points were obtained while the bead was rising. At least 700 beads were recorded for three distinct power values. Thus, a heat map was generated to represent the location dependence of the force calibration factor (Supplementary Figure 1), which relates the applied power to the acoustic force on the beads as previously reported [16].

The CBM functionalized beads are allowed to bind to the cellulose surface in the AFS chip for 15 min, following which the chip is flushed briefly at 3 μl/min until loose beads were removed from the field of view (FoV). At the studied CBM density of the beads, close to 100% of the beads were bound to the surface. All rupture force measurements follow the method described in [16]. The beads were tracked with a fixed ROI and a frame rate of 30 Hz. For the first 60 seconds, the beads were tracked in the absence of any force to determine the anchor point. Then, a linear force ramp of the required loading rate was applied until all the beads ruptured or 100% power value was reached. Each rupture force experiment was carried out in at least duplicate, with more than 60 beads tracked in each experiment.

Bead traces were analyzed using the GUI developed in [16]. After removing outliers based on a standard deviation of > 2 μm in z during the anchor point determination, each z-trace of the remaining beads was averaged using a moving window of 8-180 frames, depending on the applied loading rate. The GUI automatically determines the rupture point based on a user-defined z-value cut-off of 2 μm. The beads with tracking errors were flagged by the GUI and their rupture forces were determined manually. Based on the heatmap created during force calibration, the rupture force was determined by obtaining an interpolated force calibration factor for the respective bead position and multiplying it with the power value at the bead rupture time.

### Analyzing the rupture force histograms

The rupture force histograms from each experiment were analyzed in MATLAB. Beads that remained stuck to the cellulose surface, i.e., with a rupture force value of 0, were removed from the rupture force analysis. The number of such beads stuck to the surface at the end of the experiment are represented as final bound beads in **Figure 2** and **Figure 3**. Outliers that have a rupture force value of more than three scaled median absolute deviations (MAD) from the median rupture force, were removed, to yield a better description of the majority of ruptured beads. This resulted in removal of 2.4% of the total beads on an average. A log-normal distribution was fit to the resultant rupture force data and the mean of the log-normal fit is inferred to be the expected rupture force of the CBM-functionalized beads.

**Figure 2:**
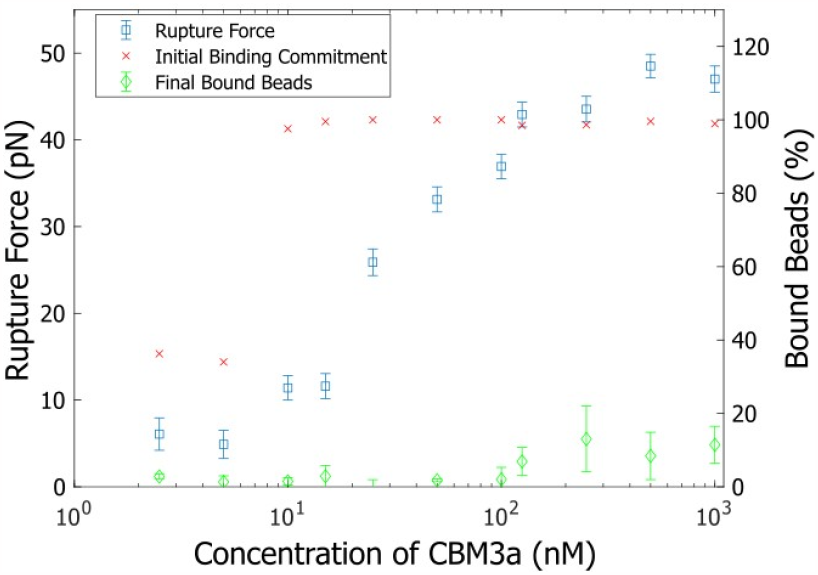
Effect of CBM concentration on the mean rupture force of CBM3a functionalized beads. The mean rupture force is obtained from the lognormal fit to the rupture force histograms and the error bars represent the standard deviation of the fit. The ratio of beads that remain bound to the surface post-flushing to the total number of beads initially in the field of view (FoV) is expressed as initial binding commitment. The percentage of final bound beads represents beads which remained bound to the cellulose surface after execution of the force ramp. The error bars of the final bound beads represent the standard deviation of the final bound bead percentages obtained during duplicate sets of experiments at each concentration.

**Figure 3:**
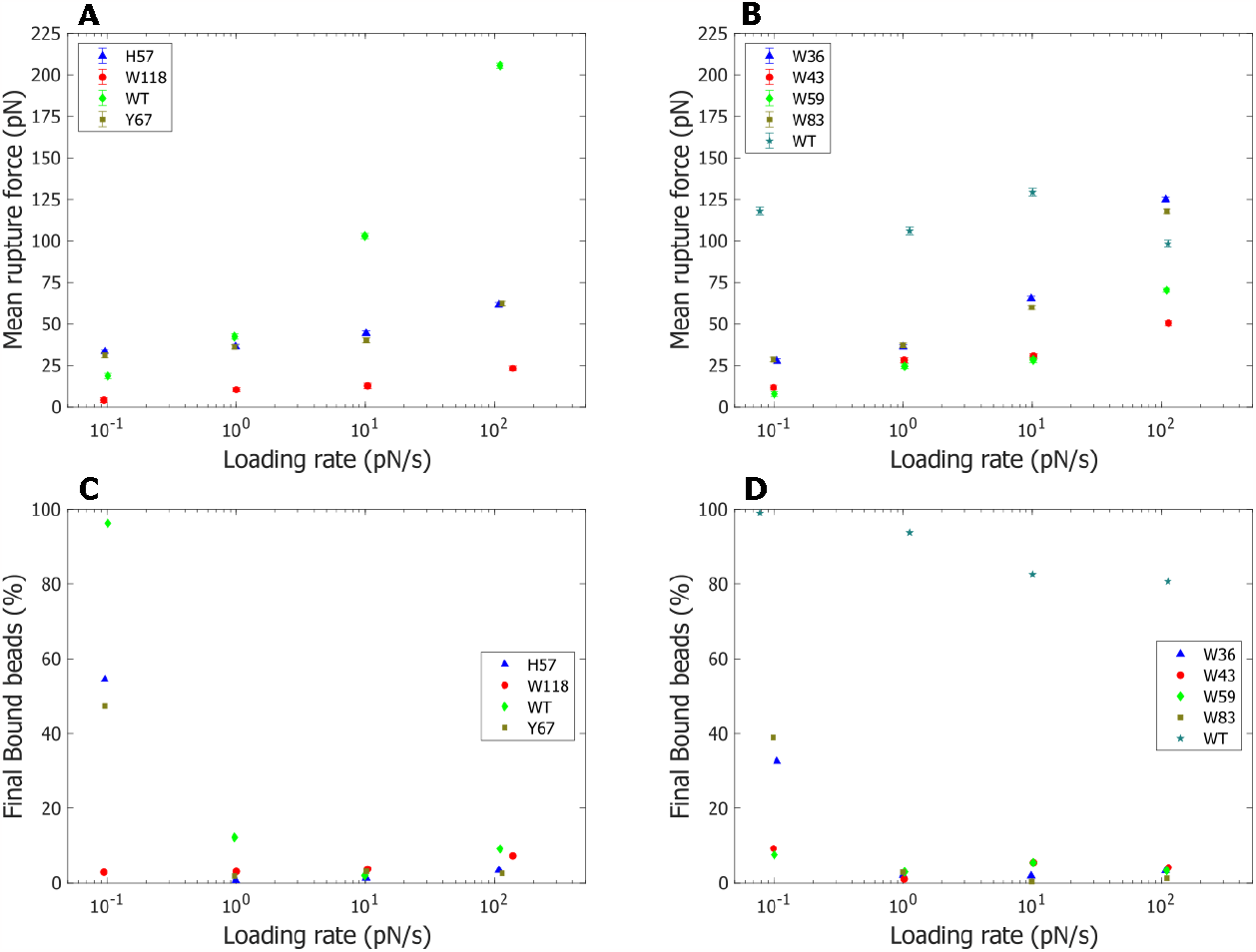
Loading rate dependence of rupture force and final bound beads for CBM3a and CBM64 wild type and mutants. The mean rupture force is obtained from the lognormal fit to the rupture force histograms of each CBM type and the error bars represent the standard deviations of the fit. The mean rupture force at various loading rates for CBM3a WT and its mutants is plotted in (A) and for CBM64 WT and its mutants in (B). The ratio of the number of beads that are stuck to the surface after completion of the force ramp to the initial number of tracked beads at various loading rates is plotted for CBM3a WT and its mutants in (C) and for CBM64 WT and its mutants in (D).

Bell-Evans model [26] is used to further quantify the bond parameters between CBM-functionalized bead and cellulose surface. Bell-Evans model, as shown in equation 1, provides the rupture force, *F as* a logarithmic function of loading rate, *r*. The fit parameters are the transition state distance, Δ*x* and bond lifetime, *k*_*off*_. *k*_*b*_ is Boltzmann constant and *T* is the temperature. The Bell-Evans model is preferred since it is relatively simple and sufficient to provide quantitative comparisons between different CBMs. Since the rupture force, *F* is that of the CBMs functionalized bead and not of single CBM-cellulose interaction, the fit parameters are referred to as *apparent k’*_*off*_ and *apparent Δx’*.

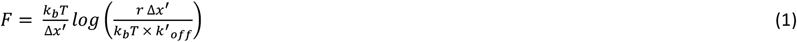

## Results and Discussion

### Characterizing the stability of the nanocellulose surface

Before each day of experiments, the rupture force of streptavidin beads, washed with WB and resuspended in 40 μl of 10 mM Phosphate buffer, was determined to verify that non-specific binding is in the expected range as an indicator for cellulose surface quality. The initial binding commitment, expressed as the ratio of beads in the FoV before and after the flushing step, was between 22% to 38% during the first 16 days of experiments (Supplementary Figure 2), compared to a near 100% binding commitment of CBM-functionalized beads. The non-specific binding is quite high when compared to the results in [16] because of lower surface passivation with the current protocol. In the same duration of time, the rupture force of streptavidin-only beads was 8.31 ± 1.52 pN (n=9) when compared to a rupture force of 46.05 ± 4.62 pN (n=5) in CBM3a WT. The sample size n here is the number of replicates over the course of 16 days and not the number of individual beads. This shows a quantitative distinction between specific and non-specific binding and allows us to judge the changes in the cellulose surface without removing the AFS chip from the instrument.

### Influence of CBM concentration on the rupture force and initial binding commitment

Since we made slight modifications to the experimental setup, we repeated experiments that characterize the rupture force of CBM3a wild type as a function of the CBM concentration [16], with the results shown in **Figure 2**. At low CBM concentrations, the rupture forces of the beads are nearly the same (compared to **Figure 2** in [16]). This is indicative that the individual CBM–cellulose interactions are not affected with the current protocol. However, the rupture force values are larger at higher concentrations, increasing at 0.28 pN/nM on an average between 10 nM and 100 nM, as compared to a 0.13 pN/nM increase in the same range in [16]. This is possibly because the CBMs have a higher availability of binding sites on the cellulose surface, where the bead is attached to, as the extent of surface passivation is much lower in the current methodology. However, the trend of rupture force versus concentration is the same as previously reported and the rupture forces appear to level off at concentrations greater than 100 nM. The initial binding commitment remains above 98% for higher concentrations but drops significantly below 10 nM.

### Effect of loading rates on the rupture force

**Figure 3** shows the loading rate (mean of normal fit to loading rate histograms) dependence of the rupture force (mean of log-normal fit to rupture force histograms) of the functionalized CBM beads. Amongst CBM3a beads, W118 has the lowest rupture force at all loading rates. CBM3a wild type (WT) has a very strong dependence on the loading rate than the other mutants, which leads to a higher apparent k’_off_ predicted by the Bell-Evans model. The proximity of the mutated sites in H57 and Y67, leading to similar binding strength can explain why the two mutants have nearly the same rupture forces at all loading rates.

‘Mutations of the central tryptophan residues, W43 and W59 of CBM64 to alanine results in lower rupture forces at all loading rates when compared to the same mutations in the peripheral residues, W36 and W83, indicating that the relative importance of central aromatic residues for the rupture force is higher. There is a noticeable difference in rupture forces of W43 and W59 mutants only at higher loading rates. The closeness in rupture forces could be attributed to similarity in binding conformations arising from point mutations in the central binding region. The same may be true for W36 and W83 mutants. With a maximum applied force of 400 pN, the rupture force of CBM64 WT does not show any significant dependence on the loading rate, in contrast to the CBM3a WT beads. Only a small fraction (<20%) of the CBM64 WT beads ruptured from the surface, however, the ruptured beads (2 ruptured beads at 0.1 pN/s to 48 ruptured beads at 100 pN/s) indicate no dependence on the loading rate. To fully assess the loading rate dependence of CBM64 functionalized beads, significantly higher forces are needed. This can be accomplished by either repeating the AFS experiment at more efficient resonance frequencies or increasing the bead size.

The rupture force vs. loading rate data of the CBMs are fit to a Bell-Evans model and the fit parameters are tabulated in **Error! Reference source not found**.

### Rupture force of binary combinations of mutant CBMs loaded on the same bead in equal proportion

**Table 1:** Fit parameters using the Bell model for different protein variants. CBM64 WT is excluded from this table due to the low number of ruptured beads along with the loading rate independence indicated as seen in **Figure 3**.

| Variant Name | Apparent $\Delta x'$<br>(nm) | Apparent $k'_{\text{off}}$<br>(*1000) ($\text{s}^{-1}$) |
| --- | --- | --- |
| CBM3a WT | 0.15 | 3.84 |
| CBM3a H57 | 1.04 | 0.01 |
| CBM3a Y67 | 0.99 | 0.02 |
| CBM3a W118 | 1.65 | 8.72 |
| CBM64 W36 | 0.29 | 2.47 |
| CBM64 W43 | 0.8 | 1.69 |
| CBM64 W59 | 0.49 | 7.63 |
| CBM64 W83 | 0.33 | 1.99 |

The fusion of different CBMs to the same catalytic domain is known to improve its enzymatic activity on plant biomass [9]. However, the dynamics of cellulase with multiple CBM-substrate interactions is not well understood. With this inspiration, we were interested in characterizing the rupture forces of beads functionalized with two types of mutant CBMs of the same family and in equal proportion to understand the extent of synergy between the CBMs while unbinding from the cellulose surface. The rupture forces of all binary combinations of the mutant CBMs, as shown in Supplementary Figure 4, are intermediate or close to the rupture force values of the parent CBMs. This indicates that there is not a significant synergistic effect leading to an improvement in binding strength due to different binding conformations of the CBMs. To check the individual contributions of parent CBMs to the overall rupture force of the multifunctional beads, we functionalized 5 μl of streptavidin beads with 5 μl of one parent CBM (H57 and Y67) and 5 μl of biotin-tagged GFP. The mean rupture force of H57+GFP bead was 19.06 ± 1.64 pN (compared to 22.85 ± 1.49 pN for H57+W118 bead) and 24.03 ± 1.44 pN for Y67+GFP bead (compared to 24.44 ± 1.49 pN for Y67+W118 bead), as shown in Figure 4. This indicates that the stronger binding mutant determines the rupture force when two mutants of significantly different rupture forces are functionalized on the same bead. These findings offer a rationale for designing multi-interaction domains using CBMs. The local variation of interactions with the target microenvironment will provide the multi-CBM-fused domains with newer dynamics to explore the microenvironment, which can potentially be used for diverse biotechnological applications in the near future.

**Figure 4:**
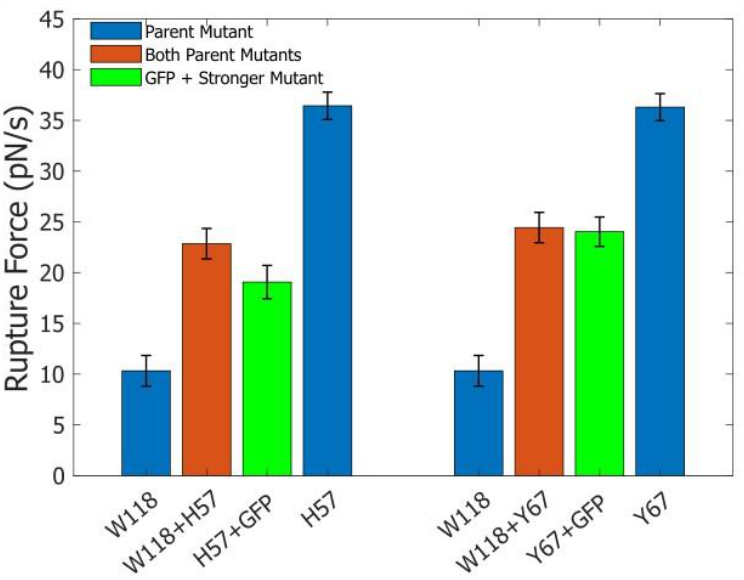
Mean rupture forces of beads functionalized with binary combinations of mutants in equal proportion, the two parent mutants and that of beads functionalized with GFP and the CBM with higher rupture force, all at a net protein concentration of 500 nM. The mean rupture force is obtained by fitting a lognormal distribution to the rupture force histograms and the error bars represent the standard deviations of the fit.

## Conclusion

We have improved an existing DFS protocol to measure the rupture forces of CBM functionalized beads and demonstrated the stability of the cellulose surface for up to 16 days, enabling the screening of a large number of mutants more conveniently. The cellulose still covered the AFS chip surface uniformly after the 16 days, as assessed by fluorescence microscopy after incubating it with GFP-tagged CBM3a WT, similar to the previously described method [17]. Though the non-specific binding of the streptavidin beads (quantified through the initial binding commitment and rupture force) is higher than previously reported [16], we have functionalized the beads with 500 nM of mutant CBMs (equimolar concentration with the streptavidin sites is reached at a concentration of 400 nM), high enough to ensure that non-specific interactions of the bead with the cellulose surface is not relevant. The rupture force data of the mutant CBMs at different loading rates can be used to estimate the transition state distance and unbinding rate using various models. Our current analysis yields apparent CBM-cellulose bond parameters, which cannot directly be compared with the other experimental data [27]. This is because the correlation between the rupture forces of functionalized beads and the number of CBMs present on them cannot be deconvoluted by our model. However, the usage of a multicomponent model [28], [29] to analyze the dissociation of CBM functionalized beads in future studies could help obtain the true single-molecule CBM-cellulose bond parameters using our experimental setup.

Finally, the insights from our experiments with beads functionalized with two different types of CBMs will be useful while designing avidity model systems with heterogeneous effectors on the same bead. It is well known that cellulosomes, possessing multiple catalytic domains and CBMs, follow a “sit and dig” mechanism [30]–[32], where the cellulosome complex degrades cellulose crystals without dissociating from the substrate. Engineering cellulosomes with CBMs having high rupture forces at some sites and CBMs having low rupture forces at other sites, will provide the designer cellulosome a strong anchorage at a particular site but allow it to explore the local environment with more flexibility. Imaging the motion of multifunctional beads in 3-D or studying the glycolytic activity of cellulosomes with multiple CBMs can further provide evidence supporting the suggested molecular motion mechanism in cellulosomes containing multiple CBMs.

## Acknowledgments

Prof. Shishir P.S. Chundawat and Mr. Gokulnath Ganesan acknowledge support from the National Science Foundation (NSF CBET Award 1846797) and Indo-U.S. Science and Technology Forum (Khorana Scholarship, 2023), respectively.

