## Supplementary Information for "Characterizing the Cellulose Binding Interactions of Type-A Carbohydrate-Binding Modules using Acoustic Force Spectroscopy"

This supplementary information includes:

Supplementary Figures 1-4

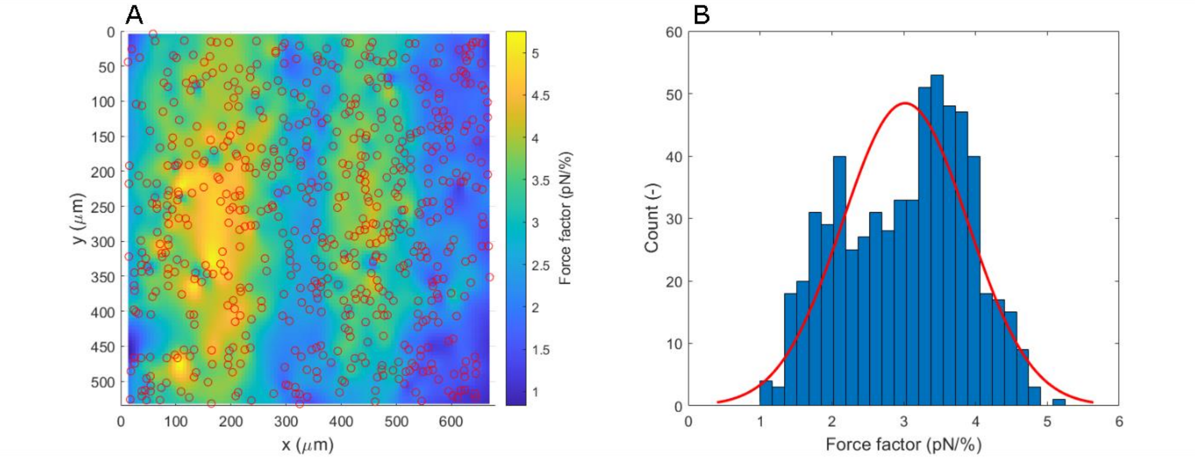


***Supplementary Figure 1: Heatmap to capture the variation of Force factor over the surface of the AFS chip.*** *(A) is a heatmap of the Force factor in the Field of View (FoV). The merged ROIs of the tracked beads during force calibration is shown as red circles. (B) is the histogram of the Force factor in the FoV, along with a normal distribution fit. The mean Force factor value obtained from the normal distribution is 3.02 ± 0.87 pN/%.*

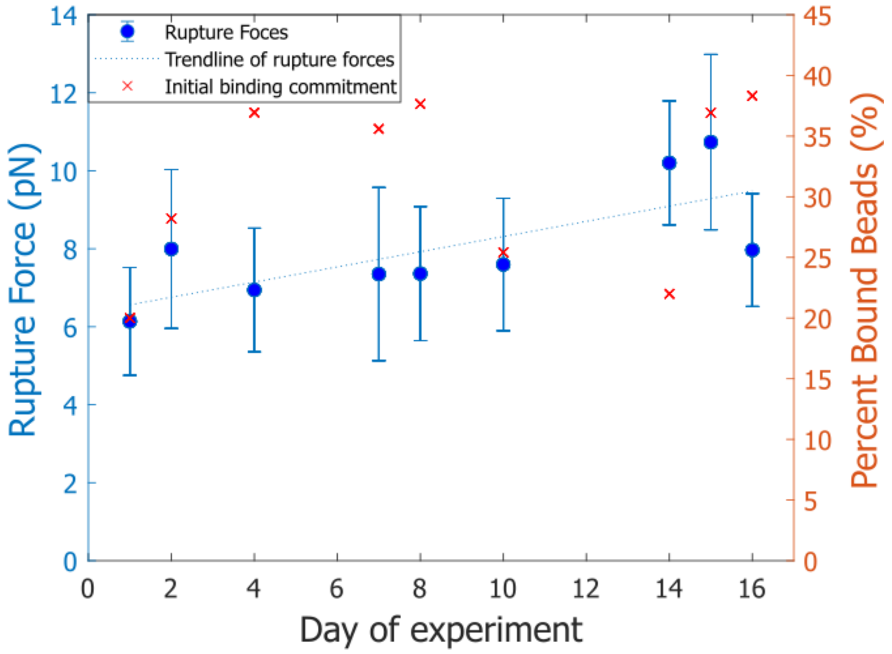


***Supplementary Figure 2: Variation of the rupture force of streptavidin beads without CBMs on days when the experiments were performed.*** *The error bars represent the standard deviation of the rupture forces of unfunctionalized beads. The rupture forces of the streptavidin beads increased by 0.19 pN/Day on average.*

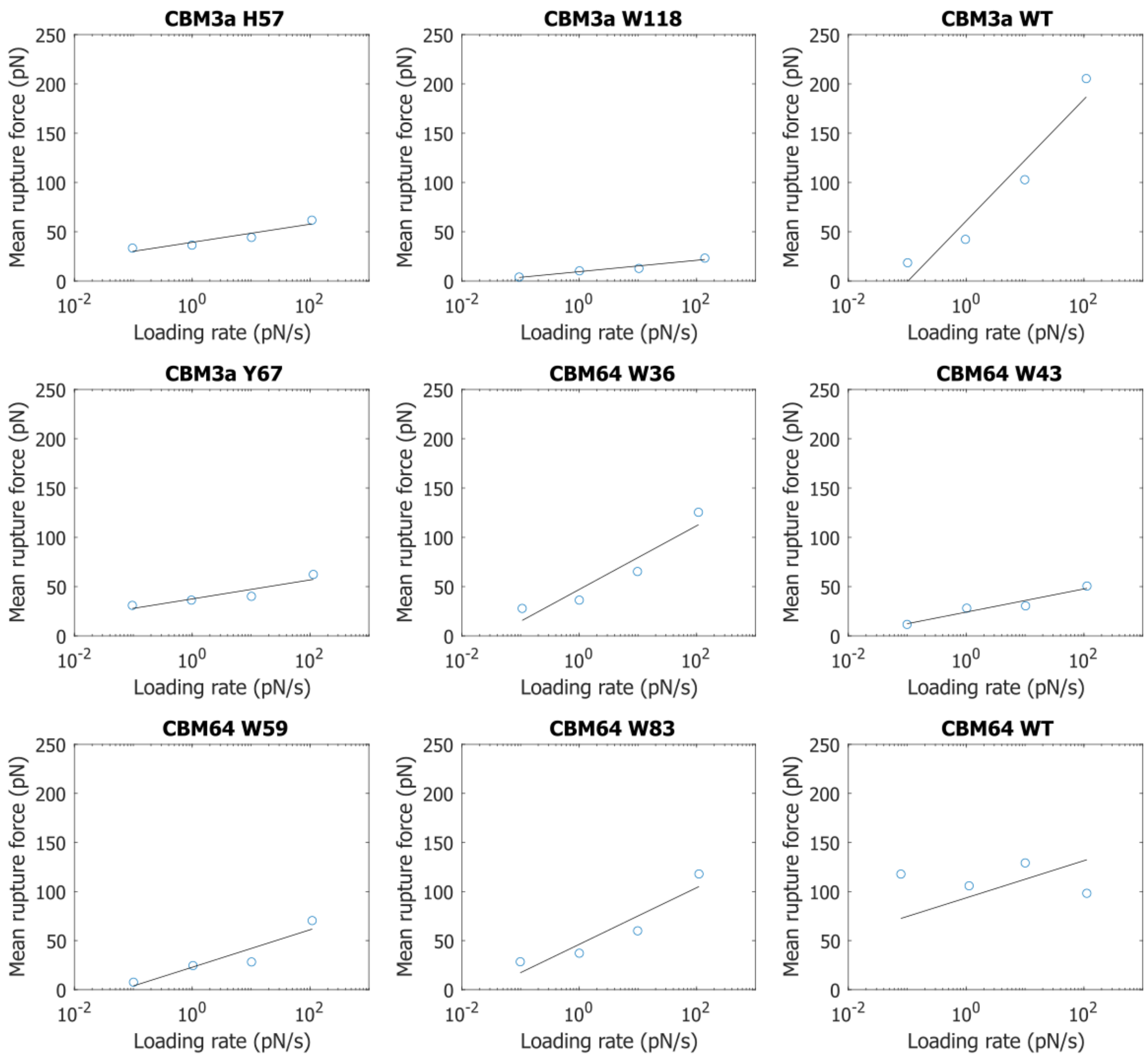


***Supplementary Figure 3: Plots of rupture force vs. loading rate, along with the Bell-Evans model fit for each mutant CBM effector bead.*** *In all cases except CBM64 WT, the Bel-Evans model fit is a good descriptor of the underlying trend between rupture forces and loading rate.*

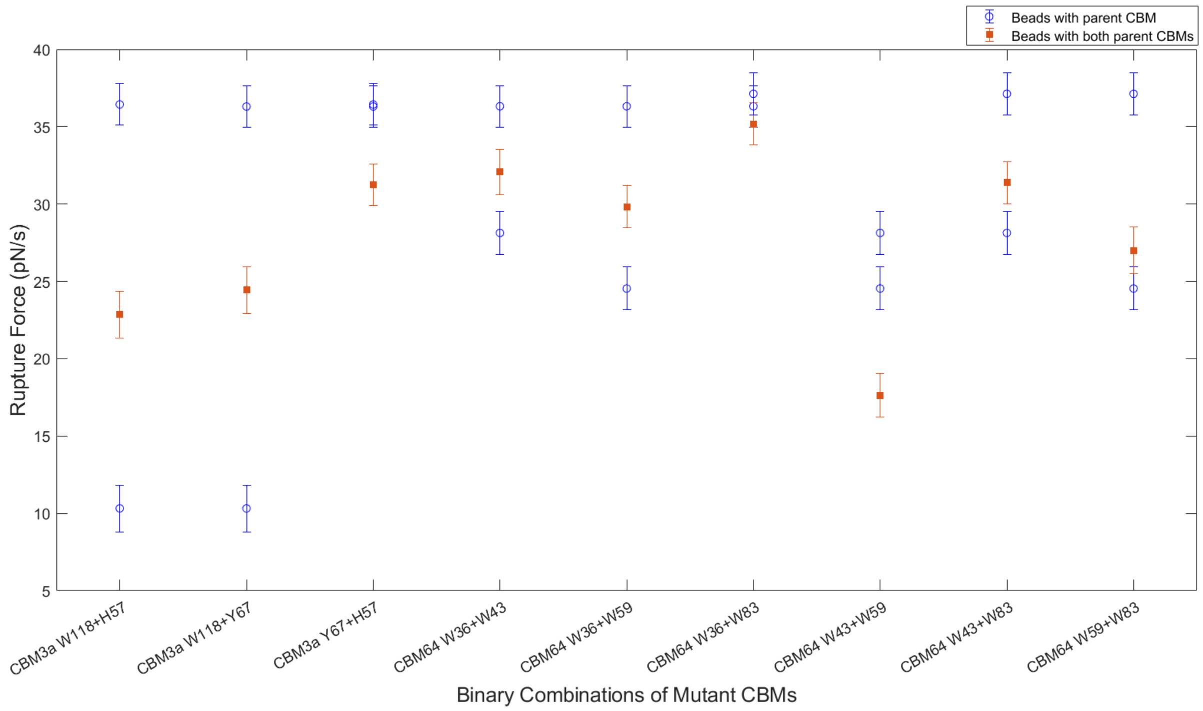


***Supplementary Figure 4: Mean Rupture forces of binary combinations of mutants loaded onto the same bead in equal proportion, along with the rupture forces of the two mutants, all at a net CBM concentration of 500 nM****. The mean rupture force is obtained by fitting a lognormal distribution to the rupture force histograms and the error bars represent the standard deviations of the fit. The rupture forces of the binary combinations Y67+H57 and W36+W83 are marginally lower than either of the parent mutants. The rupture force of W43+W59 is notably lower than both its parent mutants. This happens when the locations of mutations in both mutants are close by on the binding plane. Future molecular dynamics simulations to find the binding orientations of the proteins with respect to the cellulose surface can validate and perhaps explain this trend.*
